# Cell type-specific expression profiling sheds light on the development of a peculiar neuron, housing a complex organelle

**DOI:** 10.1101/158063

**Authors:** Kartik Sunagar, Yaara Columbus-Shenkar, Arie Fridrich, Nadya Gutkovich, Reuven Aharoni, Yehu Moran

## Abstract

Specialized neurons called cnidocytes define the phylum Cnidaria. They possess an ‘explosive’ organelle called cnidocyst that is important for prey capture and antipredator defense. An extraordinary morphological and functional complexity of the cnidocysts has inspired numerous studies to investigate their structure and development. However, the transcriptomes of the cells bearing these unique organelles are yet to be characterized, impeding our understanding of the genetic basis of their biogenesis. By generating transgenic lines of the sea anemone *Nematostella vectensis* using the CRISPR/Cas9 system, we have characterized cell-type specific transcriptomic profiles of various stages of cnidocyte maturation and show that nematogenesis (the formation of functional cnidocysts) is underpinned by dramatic shifts in the spatiotemporal gene expression. We also highlight the stark fall in transcriptional-levels of toxin and structural protein coding genes within cnidocytes with the maturation of capsule. We further reveal that the majority of upregulated genes and enriched biochemical pathways specific to cnidocytes are yet to be characterized. Finally, we unravel the recruitment of a metazoan stress-related transcription factor complex into nematogenesis and highlight its role in the formation of a structural protein of the cnidocyst wall. Thus, we provide novel insights into the biology, development, and evolution of cnidocytes.

## Introduction

Cnidocytes, also known as stinging cells, are specialized neuronal cells that typify the phylum Cnidaria (sea anemones, corals, hydroids, and jellyfish) (Kass-Simon and Scappaticci 2002; David et al. 2008; Layden et al. 2016). These cells contain an organelle called cnida or cnidocyst, which is the product of extensive Golgi secretions. Cnidae are composed of a complex capsule polymer characterized by cysteine-rich peptides, such as minicollagens and nematocyst outer wall antigen (NOWA) (Meier et al. 2007; David et al. 2008). Further, the capsule elongates at its end into a tubule that is made up of a polymer of peptides, including minicollagens, nematogalectins, and other structural proteins (Adamczyk et al. 2008; Hwang et al. 2010). This tubule invaginates into the capsule during maturation and remains tightly coiled until activated, such as during prey capture or defense, which results in the discharge of the cnidocyte capsule and uncoiling of the tubule. Cnidae are arguably the morphologically most complex organelles found in nature and their explosive discharge is one of the fastest biomechanical processes recorded in the animal kingdom (Nuchter et al. 2006; David et al. 2008).

Cnidocysts can be divided into three main categories: i) nematocysts, the dart-shaped cnidae with spines on hollow tubules that are used for prey piercing and venom injection; ii) spirocysts, the elastic cnidae used for prey entanglement; and iii) ptychocysts, the sticky cnidae that are used for adherence to prey. Nematocysts can be further divided into 25 subcategories based on their morphology (Kass-Simon and Scappaticci 2002). Such an extraordinary morphological and functional complexity of cnidocysts has driven several genetic and biochemical studies striving to identify the molecular mechanisms underlying the formation of this enigmatic organelle (Hwang et al. 2007; Milde et al. 2009; Balasubramanian et al. 2012). However, no study to date has uncovered the complete transcriptomic landscape of cnidocytes, impeding our understanding of the biology, development, and evolution of these complex organelles.

Here, we characterize the cnidocyte transcriptome of the sea anemone *Nematostella vectensis,* a rising model organism for the study of cnidarian biology (Technau and Steele 2011; Layden et al. 2016). Using the Clustered Regularly Interspaced Short Palindromic Repeats (CRISPR)/Cas9 system (Hsu et al. 2014; Wright et al. 2016), we genetically engineered transgenic lines that express a fluorescent protein in their cnidocytes. We then proteolytically dissociated tissues and separated the resulting fluorescent cnidocytes from the other types of cells using a Fluorescent Activated Cell Sorter (FACS). This led to the characterization and identification of thousands of genes that are differentially expressed at various stages of cnidocyte maturation in comparison to the other types of cells. Our findings shed light on the remarkable complexity and the dynamics of nematogenesis (cnidocyst biogenesis) or the differentiation of cnidocytes from their neuronal progenitor cells. This experimental design further enabled the identification of cnidocyte specific transcription factors that have originated in Cnidaria from a metazoan stress-related transcription factor complex around 500 million years ago. By genetically manipulating *Nematostella,* we demonstrate the role of these unique transcription factors in nematogenesis, particularly towards the formation of an integral capsule protein. Thus, our study provides a novel insight into the fascinating biology, development, and evolution of cnidocytes - the first venom delivery apparatus to appear in animals.

## Results

### Generating *a cnidocyte reporter line*

To generate a transgenic line that expresses a fluorescent reporter protein in its cnidocytes, we used the CRISPR/Cas9 system and knocked into the genomic locus of the minicollagen *NvNcol-3,* which is a major structural protein of the cnidocyst capsule wall and hence a well-characterized cnidocyte marker in *Nematostella* (Zenkert et al. 2011). We inserted the gene encoding mOrange2 (Shaner et al. 2004) with a C-terminal RAS-derived membrane tag (hereinafter referred to as memOrange2). We follow a similar approach that was described before in *Nematostella* (Ikmi et al. 2014), with the only exception being the absence of promoter and regulatory sequences in the donor construct (Fig. 1A). The microinjected embryos started expressing the fluorescent protein in their cnidocytes 72 hours post fertilization (hpf) (Fig. 1B). As expected of F0 *Nematostella* embryos, the transgene expression was restricted to small-to-medium patches (Renfer et al. 2010). The second generation (F1) exhibited specific and strong expression of memOrange2 in a very wide population of cnidocytes, with extremely strong expression in the tentacles of the polyp (Fig. 1B). Our utilization of the CRISPR/Cas9 system exploits the homologous recombination based DNArepair mechanism of the cell to insert the transgene into the native gene, and thus, places it under the control of its native regulatory elements. Hence, the expression pattern of the reporter gene accurately mirrored the native expression of *NvNcol-3* both at the mRNA and protein levels as demonstrated by double in situ hybridization (ISH) and immunostaining, respectively (Fig. 1C).

**Figure 1.**
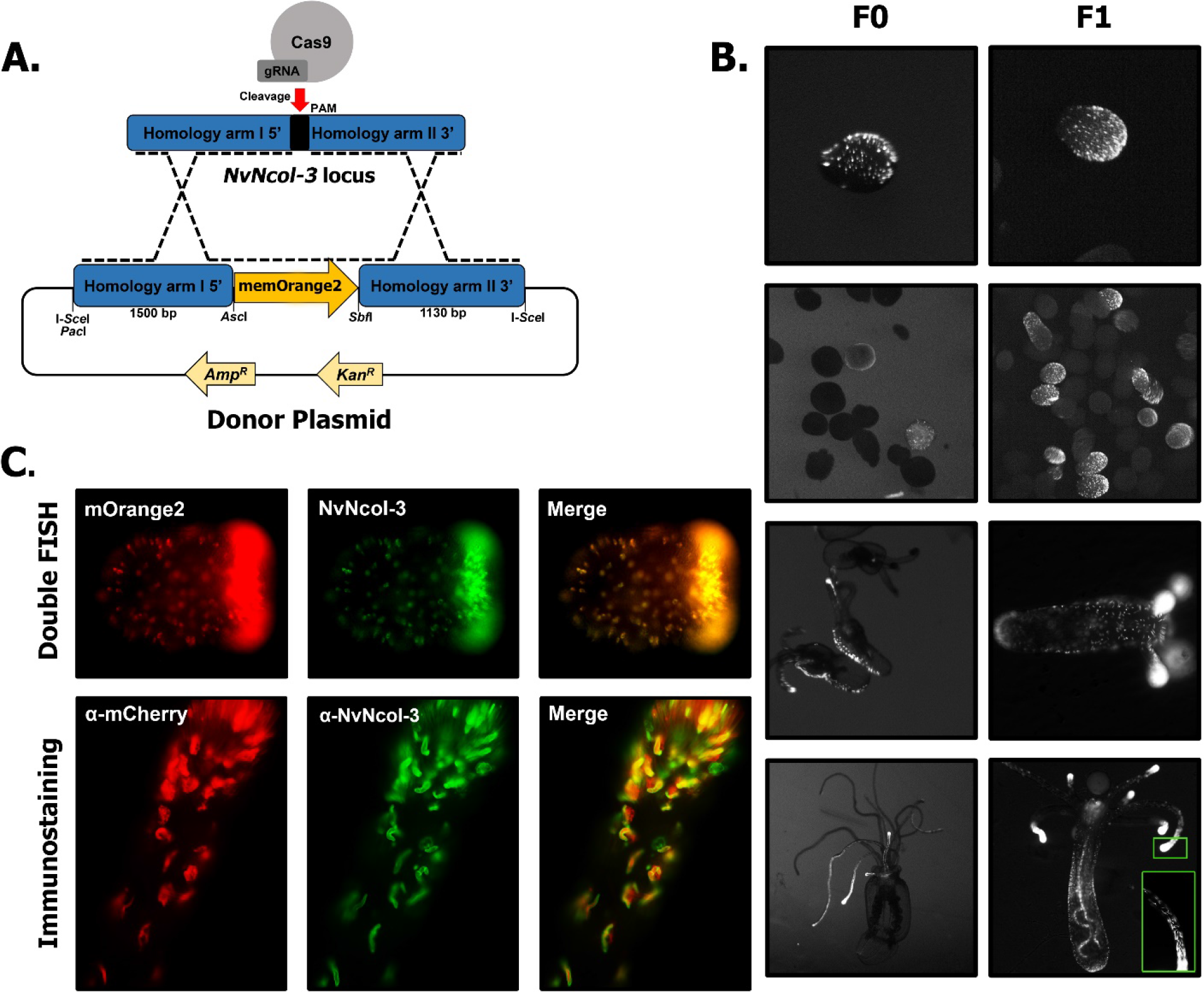
Cnidocyte reporter line. (*A*) The depiction of the reporter construct used for the generation of the transgenic reporter line. (*B*) Expression of mOrange2 in various developmental stages (planula, primary polyp, and adult) of F0 (injected) and F1 (1^st^ filial generation) are shown. (*C*) Transcriptomic and proteomic expression of *memOrange2* and *NvNcol-3,* revealed by dFISH and immunostaining, in early planulae and tentacles.

### Isolating distinct cnidocyte populations

We dissociated tentacles of ten juvenile polyps of the *NvNcol-3::memOrange2* transgenic line and sorted their cells according to the intensity of fluorescence. We initially collected two different cell populations: one that included all cells that were positive for Hoechst stain (indication of an intact nucleus) and memOrange2 (hereinafter, referred to as “positive cells”); and another population that was positive for Hoechst but negative for memOrange2 (“negative cells”) Fig. 2A-B. Then, we employed a similar strategy to collect the third population of cells that were not only positive for Hoechst but expressed very high levels of memOrange2 as well (“super-positive cells”) Fig. 2C-D. Like before, we also collected negative cells for comparison (Hoechst positive but memOrange2 negative). Microscopic inspection of fluorescent cells revealed that while the positive cells included many young developing cnidocytes, which are characterized by a round shape and small undeveloped capsules or a handful of memOrange2-filled vesicles (Supplemental Fig. 1), the super-positive cell population was largely enriched with mature cnidocytes that were easily identifiable due to the presence of elongated capsules (Fig. 2B,2D; Supplemental Fig. 1). We isolated mRNA from the aforementioned cell populations in triplicates and employed high-throughput sequencing to characterize gene expression patterns.

**Figure 2.**
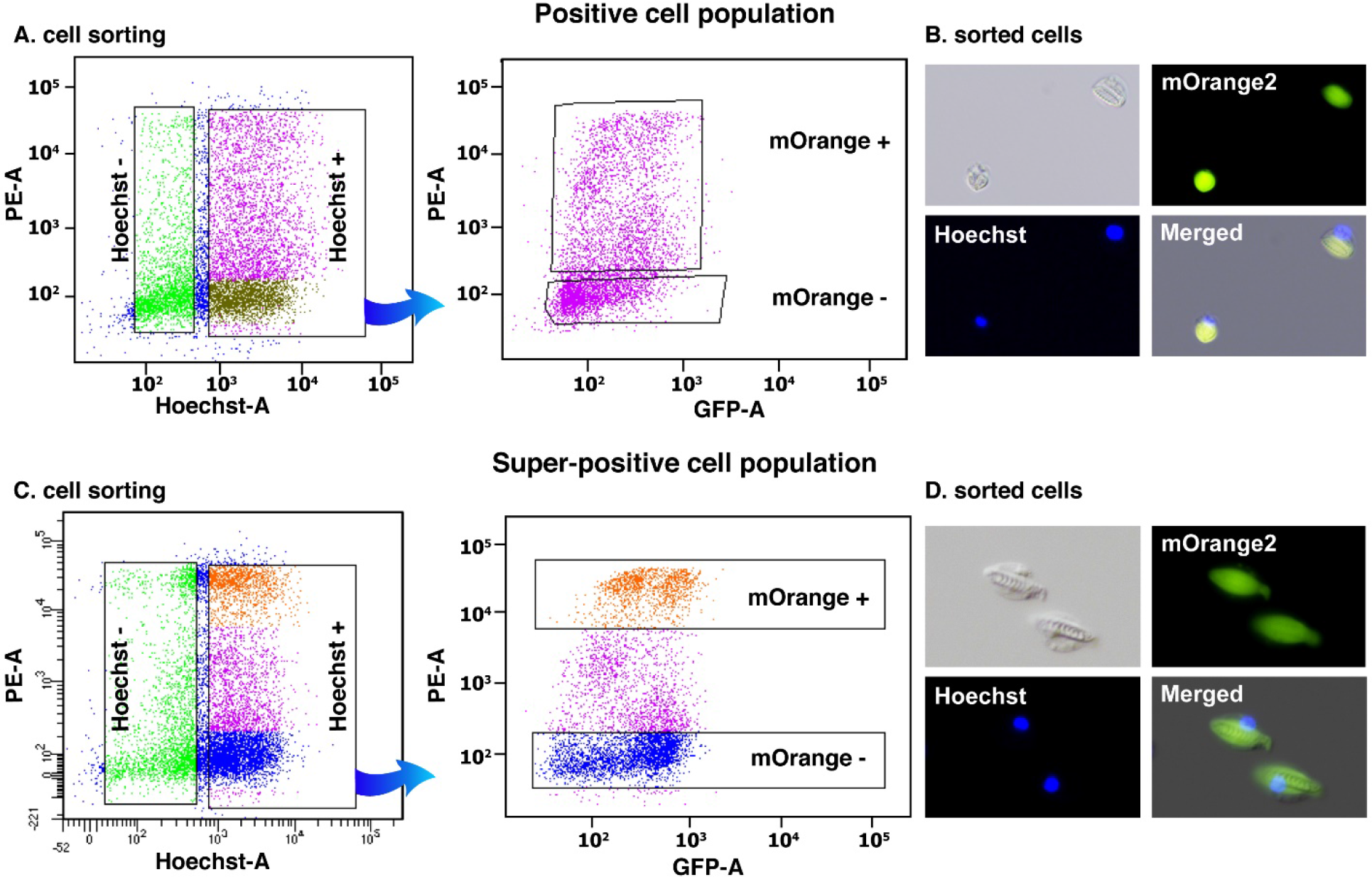
Distinct cnidocyte populations. Panels (*A*) and (*C*) highlight the FACS assisted sorting strategies employed in this study. The cells are first sorted into Hoechst negative and positive populations, followed by the separation of the latter into an memOrange2 positive, super-positive and negative populations. Panels B and D depict images of sorted positive and super-positive cells, as observed in differential interference contrast (DIC), mOrange, and DAPI filters, respectively, under a fluorescent microscope. A merger of these three images is also shown.

### Characterizing cnidocyte transcriptomes

To characterize maturation stage-specific expression profiles of cnidocytes, we employed two different comparisons: i) positives vs negatives, and ii) super-positives vs negatives. The principal component analysis revealed distinct gene expression profiles of the biological conditions (positives or super-positives vs their respective negatives) and uniform expression of biological replicates within the respective condition Fig. 3A-B. Differential expression analyses by two different methods in concert (DESeq2 and edgeR), enabled the stringent identification of a large number of differentially expressed genes in positive cell populations, in comparison to the other types of cells (Supplemental Fig. 2A-B). Our analyses revealed the significant upregulation of genes encoding known cnidocyte markers, such as *NvNcol-1, NvNcol-4, NEP-3, NEP-3-Hke, NEP-4,* and *NEP-5* (Zenkert et al. 2011; Moran et al. 2013) in positive cells (Supplemental Fig. 3A-B). Further, certain genes that are known to lack expression in cnidocytes (Marlow et al. 2009; Nakanishi et al. 2012; Wolenski et al. 2013), such as the neuronal markers *FMRFamide* and *ELAV-* the latter only identified by DESeq2, were found to be downregulated in both the positive and super-positive cells, in comparison to negative cells (Supplemental Fig. 3A-B). These results demonstrate the robustness of our experimental approach in isolating cnidocytes and accurately characterizing their transcriptional profiles. However, failing to identify cnidocyte markers as differentially expressed in the super-positive cells, despite their upregulation in positive cells, was surprising (Supplemental Fig. 3C-D). Further, the transcripts encoding memOrange2 and *NvNcol-3* were not identified as significantly upregulated in both positive and super-positive cnidocytes.

**Figure 3.**
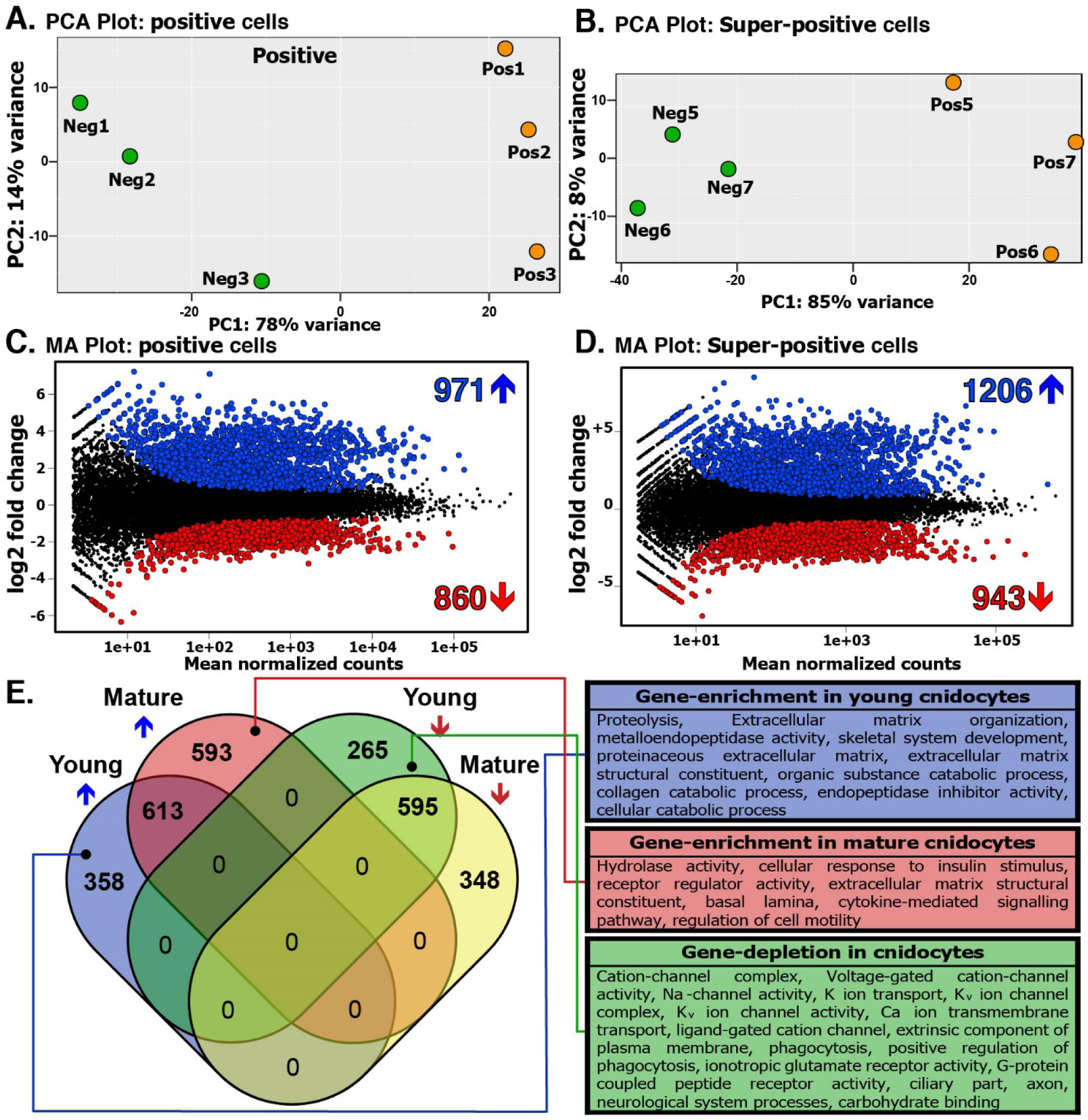
Differentially expressed genes within cnidocytes. Panels (*A*) and (*B*) show the clustering of biological replicates (memOrange2 negative: Neg1-3 and Neg5-7, positive: Pos1-3, and super-positive cell populations: Pos5-8) in a principal component analysis, where the axes represent the first two principal components, labeled with the percentages of variance associated with each axis. Panels (*C*) and (*D*) show MA-plots, which represent the log ratio of differential expression as a function of the mean intensity for each feature. The total number of upregulated and downregulated genes, highlighted in the plot as blue and red circles, respectively, are also indicated. Panel E depicts a Venn diagram, showing a comparison of differentially expressed genes identified in the positive and super-positive populations. The total number of genes uniquely identified as either upregulated or downregulated in positive and super-positive cells can be attributed to early-stage and mature cnidocytes, respectively. Enriched and depleted annotation features of each of these cell populations are also indicated.

A total of 1831 and 2149 differentially expressed genes were identified in common by DESeq2 and edgeR in positive and super-positive cell populations, respectively (Fig. 3C-D; Supplemental Table 1). Overall, 613 genes were identified as upregulated, while 595 genes were found to be downregulated in both positive and super-positive cell populations, in comparison to the negative cells (Fig. 3E). Interestingly, 358 and 265 genes were identified as significantly upregulated and downregulated, respectively in the positive cell population alone. These differentially expressed genes, identified only in the positive cells, can be attributed to the earlier stages of maturating cnidocytes, which get depleted in the super-positive population that is enriched with mature cnidocytes. Similarly, we identified 593 and 348 genes as uniquely upregulated and downregulated, respectively, in the mature cnidocyte-rich, super-positive population (Fig. 3E).

### Spatiotemporal dynamics of cnidocyte-specific transcripts and their encoded proteins

The absence of significant enrichment of *NvNcol3* and *memOrange2* transcripts in both the positive and super-positive cell populations was surprising. We suspected that this might be connected to the maturation stage of the isolated cnidocyte and that, perhaps, the temporal differences in expression levels of *memOrange2* RNA and protein may result in an inability to capture very early stage cnidocytes. In order to test this hypothesis, we performed an ISH experiment, followed by Immunostaining. These combined assays enabled us to distinguish mRNA and protein expression of *NvNcol-3* and *memOrange2in* the 3-day-old wild type (Fig. 4A) and *NvNcol-3*::*memOrange2* transgenic planulae (Fig. 4B), respectively.

**Figure 4.**
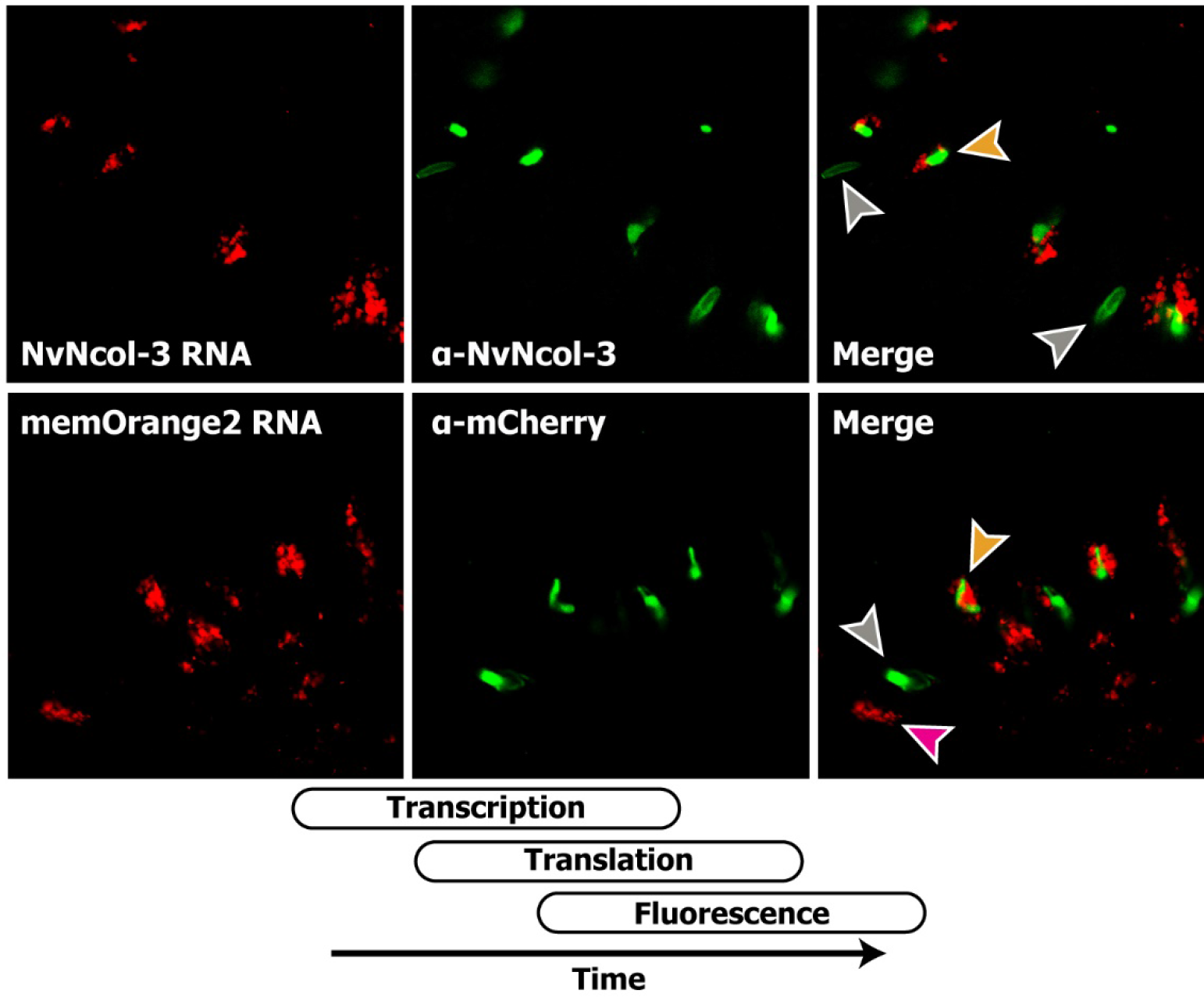
Variability between the transcriptional and proteomic expression levels of cnidocyte-expressed genes. This figure shows that the expression pattern of *NvNcol-3* and *memOrange2* genes (FastRed, red) differs from the expression pattern of their protein products (Alexa fluor 488, green). Examples for cells containing only RNA, only protein or both RNA and protein are indicated by magenta, gray and orange arrowheads, respectively. The schematic diagram shows the overlap between the times of gene expression, protein secretion and protein maturation.

This experiment revealed dramatic temporal differences in the mRNA and protein expression patterns of both these genes within cnidocytes. In both the wild type and *NvNcol-3* transgenic planulae, a large proportion of cnidocytes exclusively stained either at the mRNA level or at the protein level Fig. 4A-B. The cells that only expressed RNA were round and lacked capsules or contained undeveloped capsules and multiple vesicles, and were identified as cnidoblasts or early-stage cnidocytes (Fig. 4A-B; Supplemental Fig. 1), whereas those containing only protein, housed fully developed or nearly-mature capsules (Fig. 4A-B; Supplemental Fig. 1). Only a smaller fraction of the cnidocytes, most of which were characterized by relatively small and immature capsules – typical of developing cnidocytes, exhibited expression at both RNA and protein-levels Fig. 4A-B.

### The discovery of novel genes with cnidocyte-specific expression

One of the major objectives of this study was to identify novel genes with cnidocyte-specific expression, as this would advance our knowledge regarding the biology, development, and the evolutionary origin of cnidocytes. First, to test the accuracy of our experimental and bioinformatic approaches for identifying differentially expressed genes in cnidocytes, we chose nine genes that were found to be significantly upregulated in positive cells (foldchange in the range of 8 to 47 with at least 50 mapped reads in the negative sample) (Fig. 5). Interestingly, none of these nine genes were previously described in the cnidocytes of *N. vectensis*. These genes encode proteins that show a significant sequence similarity to three protein disulfide isomerases (*NVE15732, NVE15733,* and *NVE26200*), a homolog of the c-Jun transcription factor (*NVE16876*) that we named “Cnido-Jun”, a highly derived homolog of NOWA (*NVE17236*), an M13 peptidase (*NVE13546*), a calmodulin (*NVE22513*), an unknown protein containing thrombospondin and F5/8 type C domains (*NVE5730*), and a protein containing a galactose binding lectin domain with an apparent homology to Nematogalectin (*NVE3843).* The localization of expression using ISH highlighted the cnidocyte-specific expression of all of these nine genes (Fig. 5). Unexpectedly, the expression pattern of individual genes exhibited a dramatic spatiotemporal variability (Fig. 5). For example, the expression of the NOWA-like gene was limited to very few cnidocytes in the oral region of the planula larva and the tentacles of the primary polyp, while the expression of Cnido-Jun was localized to both the oral and the central part of the planula, followed by an increased expression in the tentacles of the primary polyp (Fig. 5). The three protein disulfide isomerases also showed noticeably distinct expression patterns, strongly indicating functional specialization of these enzymes (Fig. 5).

**Figure 5.**
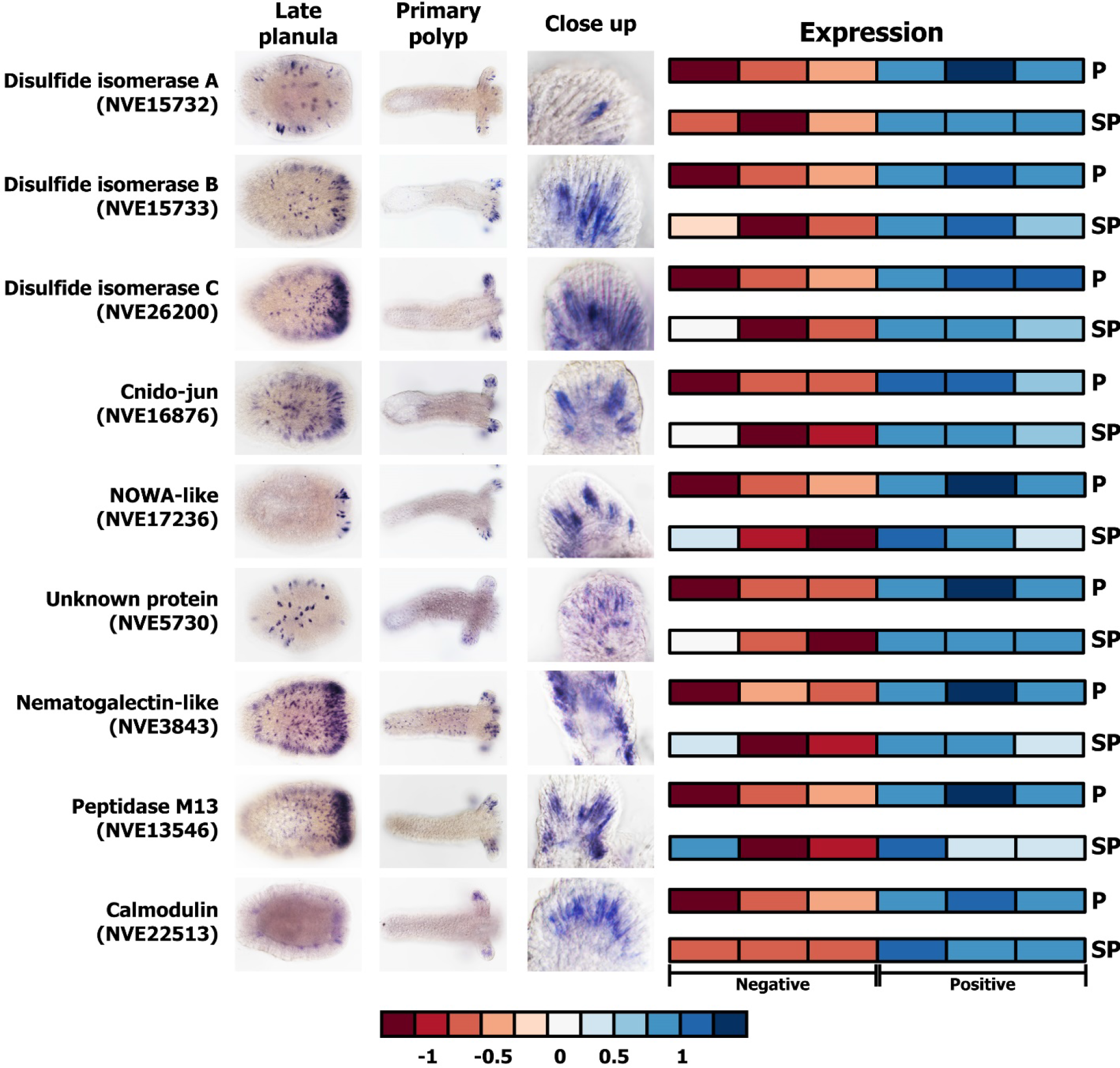
Novel genes with cnidocyte-specific expression. In situ hybridization expression patterns of novel cnidocyte-specific genes identified in this study are shown in late planula and primary polyp. The elongated cells in the ectoderm, seen in the close up image, can clearly be identified as cnidocytes according to the large unstained space, which is the cnidocyst capsule. A heatmap of expression in positive (P) and super-positive (SP) cell populations, across the technical replicates, are also indicated. A color code for expression values, ranging from a gradient of red (downregulated) to blue (upregulated), is also provided.

### Functional annotation of biochemical pathways associated with cnidocytes

Functional annotation of differentially expressed genes revealed that a large number of upregulated genes in cnidocytes (46% and 36% in positive and super-positive samples, respectively) are taxonomically restricted. Further, gene enrichment analyses enabled the identification of functional categories that are significantly enriched or depleted in positive and super-positive cell populations, providing novel insights into the biochemical pathways of these enigmatic cells (Fig. 3; Supplemental Fig. 4). The strong enrichment of hydrolytic activity in the positive and super-positive cells was identified. The enrichment of the terms related to the extracellular matrix (Fig. 3; Supplemental Fig. 4) supports a strong evolutionary link between the structural components of the cnidocyst and the constituents of the extracellular matrix (Ozbek 2011). Surprisingly, despite cnidocytes being specialized neurons known for expressing several ion channel subtypes (Bouchard et al. 2006; Moran and Zakon 2014; Li et al. 2015), a depletion of genes associated with ion channels, such as “cation transport”, “sodium-”, and “potassium ion channels” was found (Fig. 3; Supplemental Fig. 4).

### The recruitment of a stress-response transcription factor complex into nematogenesis

By retrieving homologues from the genomes and transcriptomes of a diversity of cnidarian species, we reconstructed the phylogenetic histories of two transcription factors that were found to be specifically upregulated in cnidocytes (*NVE16876 and NVE5133;* Fig. 6). We named these transcription factors Cnido-Jun (17- and 3-fold upregulation in positive and super-positive cells, respectively) and Cnido-Fos1 (33- and 20-fold upregulation in positive and super-positive cells, respectively) to distinguish them from their paralogues encoding proto-oncogenic transcription factors c-Jun and c-Fos. These transcription factors are known to dimerize into an Activation protein-1 (AP-1) complex, which is involved in various stress responses in Bilateria and Cnidaria (Hess et al. 2004; Meng and Xia 2011; Elran et al. 2014, Agron et al. 2017). We show that Cnido-Jun and Cnido-Fos1 protein coding genes originated via gene duplication in the common ancestor of Hexacorallians – sea anemone and stony corals Fig. 6A-B. However, only in sea anemones, the c-Fos gene underwent an additional round of duplication and led to the origination of Cnido-Fos2 (*NVE23145*), which was not identified as differentially expressed gene. Domain scanning of these transcription factors revealed cnidarian-specific insertions between the Jun and bZIP domains – the latter is required for DNA binding in the c-Jun proteins (Fig. 6). In a complete contrast, we found insertions in bilaterian c-Fos proteins that were missing in their cnidarian counterparts. Interestingly, Cnido-Fos1 was the only protein that had different residues than those implicated in the heterodimer formation (Fig. 6; Supplemental Fig. 5), which may suggest that this derived protein binds to novel partner proteins.

**Figure 6.**
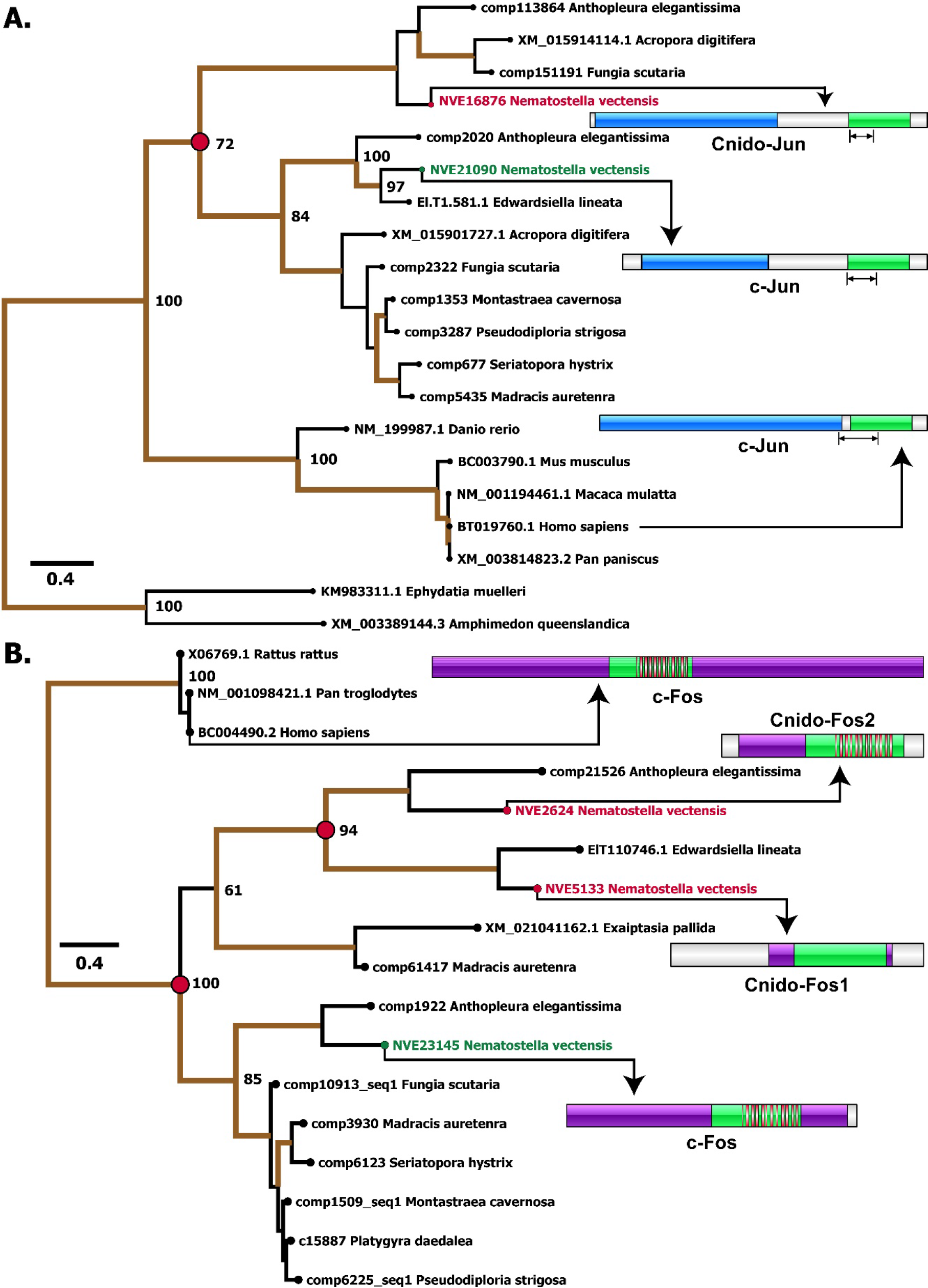
The deep origin of a novel cnidocyte-specific transcription factors in Cnidaria. This figure depicts the phylogenetic histories of the (*A*) c-Jun and (*B*) c-Fos family of proteins in Cnidaria. The duplication events that led to the origin of Cnido-Jun and Cnido-Fos1, cnidocyte-specific transcription factors identified in this study (labelled in red font), from c-Jun and c-Fos (indicated in green font) in Hexacorallia is shown in a red circle. Branches with node-support ≥ 70 are shown in thick brown lines, and the bootstrap values for the major nodes are indicated. Domain organization of these proteins are also depicted, where the Jun-like domain, the bZIP domains and the c-Fos related domains are indicated in blue, green, and purple, respectively. The DNA binding domain is marked by black arrows, while the dimerization domains are depicted in red bars.

To understand whether Cnido-Jun, which exhibit cnidocyte-specific expression, play a significant role on nematogenesis, we genetically manipulated *N. vectensis* embryos. Using the CRISPR/cas9 system, we knocked into the *NvNcol-3locus* a modified version of the Cnido-Jun missing the bZIP-domain, which ensures that it cannot function as a transcription factor (Fig. 7A). In addition to this deletion, the transgene included a sequence that encodes memOrange2, a self-cleaving P2A peptide that separates memOrange2 sequence from the manipulated Cnido-Jun sequence upon translation (Kim et al. 2011), and a c-myc Nuclear Localization Signal (NLS) (Dang and Lee 1988) (Fig. 7A). Overexpression under the native *NvNcol-3* regulatory region enables the dimerization of the altered Cnido-Jun with its native partner protein (possibly Cnido-Fos), preventing the latter from performing its biological function in cnidocytes. The genetically-manipulated anemones exhibited a mosaic expression pattern of the synthetic gene as expected of F0 animals. Strikingly, NvNcol-3 expression was completely absent from the transgenic patches that overexpress the altered Cnido-Jun protein in cnidocytes (Fig. 7B).

**Figure 7.**
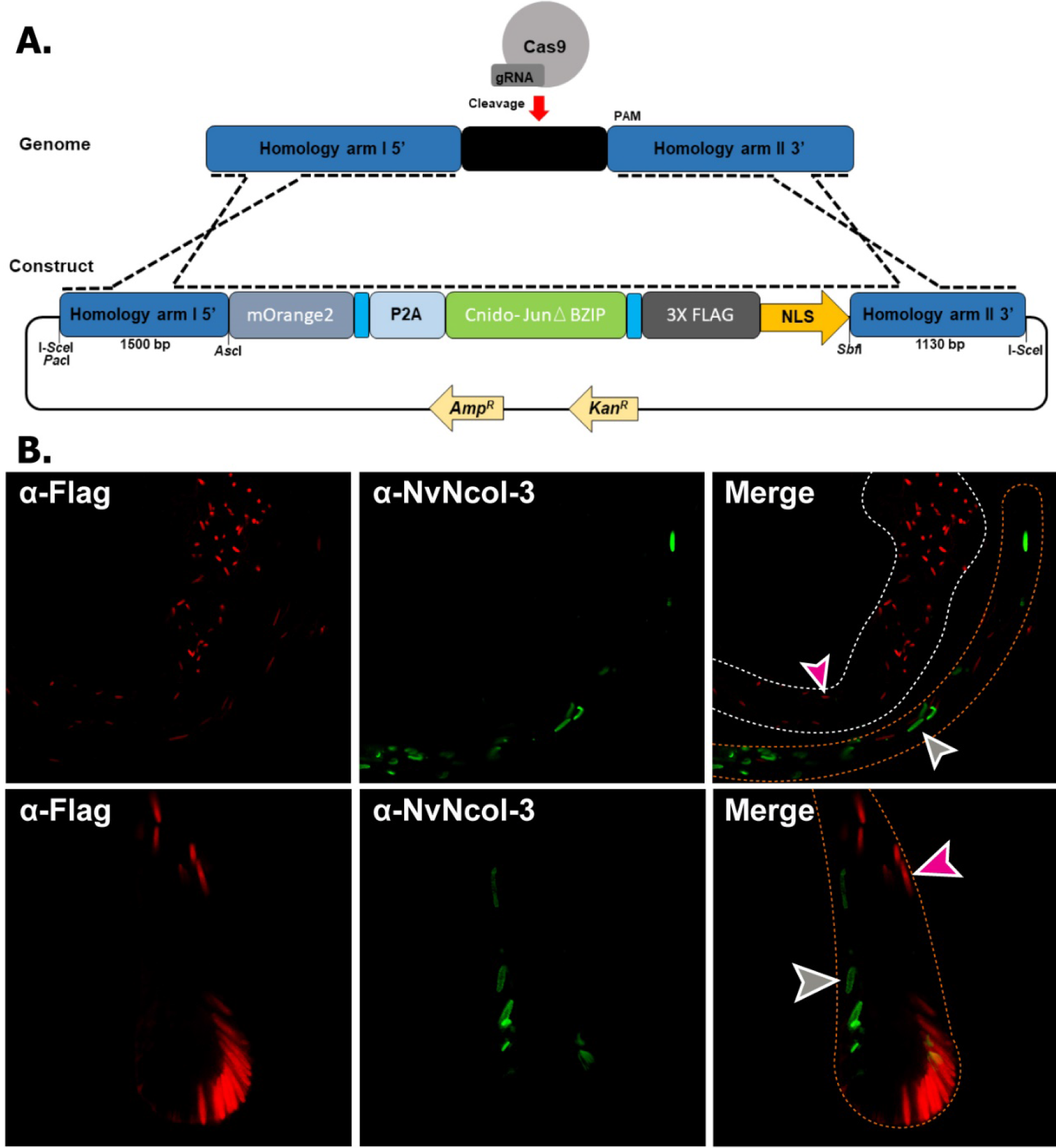
Overexpression of a truncated Cnido-Jun mutant depletes NvNcol-3. (*A*) The depiction of the reporter construct used for overexpressing the truncated Cnido-Jun mutant. (*B*) The complete lack of co-localization between the Cnido-Jun mutant and NvNcol-3 as revealed by immunostaining with α-FLAG and α-NvNcol-3, respectively, suggests the depletion of NvNcol-3 by the overexpression of the mutant. The magenta arrowhead points to a FLAG-positive cell that lacks any NvNcol-3 expression, while the gray arrowhead points to an opposite example. The body column and the tentacles are highlighted in white and orange dotted lines, respectively.

## Discussion

### A complex expression landscape across developing cnidocytes

Among the large number of upregulated genes detected within the positive cell population (Fig. 3C-D; Supplemental Fig. 2A-B), many were cnidocyte-marker genes, including toxin (*NEP-3*, *NEP-3-like*, *NEP-4,* and *NEP-5*) and structural protein coding genes (*NvNcol-1* and *NvNcoi-4*). At the same time, there was downregulation of certain neuronal marker genes (*FMRFamide* and *ELAV*) that are known to lack expression in cnidocytes (Supplemental Table 1; Supplemental Fig. 3A-B). Though there was downregulation of *FMRFamide* and *ELA V* we did not identify the upregulation of cnidocyte-marker genes in the super-positive cell population, which was enriched with mature cnidocytes. This can be explained by the fact that in mature cnidocytes the capsule, which is a very tight polymer of various peptides and proteoglycans, is already formed and the secretion of structural proteins is no longer required and is most probably wasteful. As a result, the expression of such genes diminishes with the maturation of cnidocytes. This is clearly evident when examining the expression of memOrange2 across various cell types. In our reporter line, memOrange2 was integrated into the genomic locus of *NvNcol-3*–the gene coding a vital structural protein of the capsule (Fig. 1A). The expression of memOrange2 in the positive cell population was relatively higher than the negative cell population, but it dropped significantly in the super-positive population, where it was identified as a highly downregulated gene (-1.67 log_2_ fold change; p-value: 0.0005) (Supplemental Fig. 3). This clearly demonstrates that the expression of structural proteins drops significantly with the maturation of capsules in fully-developed cnidocytes. Since the development of the mature capsule impedes the migration of secreted toxins, a similar pattern of expression can be expected for toxin coding genes at this stage of development. As explained in the following section, these results can also be attributed to the inherent dynamics of gene expression across developing cnidocytes.

It should be noted that because the secreted memOrange2 protein requires four and a half hours at 37^o^C for t_1/2_ maturation (Shaner et al. 2008), a smaller fraction of cnidocytes in the very earlier stages of development may contain an immature non-glowing memOrange2, and therefore, would get sorted into the negative population. As a result, despite the increased expression in positive cells in comparison to the negative cells, memOrange2 was not identified as a differentially expressed transcript. As explained in the following section, the temporal variation in transcription and translation of memOrange2 could also contribute to this effect. However, the robustness of our experimental approach in isolating cnidocytes for generating cell-type specific transcriptomic profiles is strongly supported by multiple lines of evidence: i) the significant upregulation of cnidocyte-specific markers (ranging between 4 to 25 fold change difference) and the downregulation of neuronal markers in positive cells; ii) microscopic examinations of isolated cells; iii) in situ hybridization experiments for the nine highly upregulated genes in cnidocytes (ranging between 8 to 47 fold change difference); and iv) the functional annotation of upregulated transcripts. Moreover, the microscopic examination of the positive cell populations revealed many cnidocytes in a very early stage of maturation (Supplemental Fig. 1). These cells can clearly be seen to contain undeveloped capsules and multiple secretory vesicles carrying premature cnidocyst components, which indicate early stages of nematogenesis when the capsule starts to form by massive Golgi secretions (David et al. 2008). Many were even completely missing capsules but contained only memOrange2-filled vesicles (Supplemental Fig. 1), suggesting that our approach can capture cnidocytes in very early stages of maturation.

### Unexpected differences in the spatiotemporai expression of cnidocyte-specific transcripts and their encoded proteins

The dramatic difference we discovered in the expression profiles of certain genes in positive and super-positive cell populations can be further attributed to the inherent dynamics of transcription and translation across the various stages of cnidocyte maturation. This was revealed by a combination of ISH and dFISH experiments, where we colocalized the transcripts and proteins encoded by *memOrange2* or *NvNcol-3* genes. With this approach, we detected three different populations of cnidocytes: i) cnidocytes in the very early stages of maturation that only contained abundant transcripts but no proteins, suggesting that they were yet to develop a capsule; ii) cnidocytes with high-levels of both transcripts and proteins, suggesting that these were developing cells but contained an immature capsule; and iii) mature cnidocytes with fully developed capsules that only contained proteins and completely lacked transcripts for *memOrange2* and *NvNcol-3* genes (Fig. 4). This supports our hypothesis that transcription of toxin and structural protein coding genes drastically falls in mature cnidocytes with the complete formation of the tightly polymerized capsule. Thus, we reveal dramatic spatiotemporal dynamics of transcription and translation of certain genes within cnidocytes (Fig. 4).

### The discovery of novel cnidocyte-specific transcripts

Our experimental and bioinformatic approaches facilitated the identification of a large number of cnidocyte-specific genes, many of which were taxonomically restricted (46% and 36% in positive and super-positive samples, respectively; Supplemental file 1), highlighting a paucity of knowledge regarding the biology of cnidocytes. We identified nine genes that were significantly upregulated in positive cells (foldchange in the range of 8 to 47) and were not known to be specifically expressed in cnidocytes. The localization of expression in ISH experiments revealed that all of these nine genes exhibited cnidocyte-specific expression (Fig. 5), albeit with a dramatic spatiotemporal variation. For example, three of these genes encoded protein disulfide isomerases (*NVE15732, NVE15733,* and *NVE26200*), which probably mediate the folding of toxins and structural proteins within cnidocytes, as both of these groups of proteins usually require the formation multiple disulfide bridges for carrying out their function (Mouhat et al. 2004; David et al. 2008; Ozbek 2011). Each of these enzymes exhibited distinct expression profiles (Fig. 5), which is suggestive of functional specialization within different types of cnidocytes. Similarly, the expression of the NOWA-like gene was limited to a smaller population of cnidocytes in the oral region of the planula and the tentacles of the primary polyp. In *Hydra,* NOWA has been implicated in the formation of nematocyst (Engel et al. 2002). If this protein serves a similar structural function in the cnidocytes of *Nematostella,* then the documented differences in expression profiles are suggestive of distinct structural makeup of different cnidocyte populations.

Interestingly, the expression of a calmodulin (*NVE22513*), a highly-conserved protein known to bind Ca^2+^ ions (Friedberg and Rhoads 2001), was restricted only to certain regions of the tentacles in the primary polyp (Fig. 5). Fluids of the mature cnidocytes are known to contain high concentrations (ca. 500-600 mmol/kg wet weight) of Ca^2+^ ions that are bound by certain proteins in the undischarged state (Lubbock et al. 1981), and the dissociation of Ca^2+^ ions from these partner proteins has been implicated in the rapid discharge of cnidocysts. Thus, the restricted expression profile of calmodulin points to the functional specialization of Ca^2+^ ion binding proteins in different types of cnidocytes. Thus, the identification of numerous novel cnidocyte-specific genes has advanced our knowledge regarding the biology and development of cnidocytes.

### The deep origin of cnidocyte-specific transcription factors from a bilaterian stress-response complex and their significance in nematogensis

Our analyses enabled the discovery of Cnido-Jun and Cnido-Fos1, two novel cnidocyte-specific transcription factors with a sequence similarity to the c-Jun and c-Fos family of proteins. c-Jun and c-Fos are known to constitute the AP-1 early-response transcription factors that are associated with stress, infection, cytokines, and other stimuli (Hess et al. 2004;). In *Nematostella,* one such c-Jun protein family member (*NVE21090*)has been identified as a player in stress response (Elran et al. 2014, Agron et al. 2017). Results of our phylogenetic analyses revealed that Cnido-Jun and Cnido-Fos1 proteins originated via duplication events in the common ancestor of hexacorallians (sea anemones and corals), approximately 500 MYA (Shinzato et al. 2011) (Fig. 6). These two genes were found to be significantly upregulated within cnidocytes, ranging between 17-33 fold upregulation, in comparison to the other types of cells. However, unlike Cnido-Jun and Cnido-Fos1, the expression of c-Jun and c-Fos stress response transcription factors was not upregulated in cnidocytes when compared to other cells. Therefore, it is very likely that the cnidocyte-specific Cnido-Jun and Cnido-Fos1 proteins, which might dimerize to form a ‘Cnido-Ap1’ complex within cnidocytes, have been recruited into a function other than stress response.

To understand the role of these unique transcription factors within cnidocytes, we genetically manipulated *Nematostella* embryos to overexpress an altered version of Cnido-Jun specifically within cnidocytes. This mutant protein lacked DNA binding domains but retained its native dimerization domains. Thus, the overexpression of this mutant Cnido-Jun prevents the normal functioning of its partner protein, and hence, and hence, the transcription factor complex itself. Surprisingly, when we co-localized the protein product of this transgene along with the native *NvNcol3* in an immunohistochemistry experiment, we discovered that the cnidocytes that express the Cnido-Jun transgene almost never expressed the native Nv-NCol-3 (Fig. 7B). This result suggests that *NvNcol3,* a gene encoding one of the better-characterized nematocyst capsule components, is either a direct or indirect target of Cnido-jun and its partner proteins.

### Functional annotation reveals differential enrichment and depletion of biochemical pathways in cnidocytes

The functional annotation of upregulated genes in positive and super-positive cell populations supported the earlier findings in *Hydra* showing that a significantly large proportion of cnidocyte-specific genes in Cnidaria are taxonomically restricted (Milde et al. 2009). Gene enrichment analyses facilitated the discovery of significantly enriched or depleted annotation categories in the positive and super-positive cell populations, providing the first insights into the biochemical pathways of these peculiar cell types (Fig. 3; Supplemental Fig. 4). A strong enrichment of annotations related to the hydrolytic activity was noted in the positive and super-positive cells, which can be explained by the abundant presence of proteolytic and hydrolytic enzymes in cnidocytes that are responsible for processing cnidocyst structural components, as well as for exerting toxicity in prey or predatory animals (Balasubramanian et al. 2012; Moran et al. 2013; Rachamim et al. 2015). Annotation terms related to the extracellular matrix, such as the “basal lamina” (Fig. 3; Supplemental Fig. 4), were found to be enriched in positive and superpositive cells. This supports the proposed evolutionary link between cnidocyst structural components and the extracellular matrix constituents (Ozbek 2011). Interestingly, despite the fact that cnidocytes are specialized neurons that are known to express several subtypes of ion channels (Bouchard et al. 2006; Moran and Zakon 2014; Li et al. 2015), a depletion of annotation terms related to ion channels was identified (Fig. 3; Supplemental Fig. 4). This can be explained by the fact that many subtypes of ion channels are known to be specifically expressed in certain other cell types, such as neurons and muscles, but not in cnidocytes (Gur Barzilai et al. 2012; Jegla et al. 2012; Moran and Zakon 2014; Grunder and Assmann 2015). The surprising enrichment of annotation terms like “cytokine-mediated signaling pathway”, “cellular response to insulin stimulus”, and “regulation of cell motility” in cnidocytes, further highlights the presence of several uncharacterized biochemical pathways that might be relevant to cnidocytes. Thus, several specialized ion channels and biochemical pathways appear to be unique to cnidocytes.

We conclude that nematogenesis or the differentiation of neuronal precursor cells into cnidocytes, is a highly dynamic and multistep process that is accompanied by complex shifts in the transcriptional and translational profiles of differentiating cnidocytes. We highlight a drastic reduction in the transcriptional profiles of toxins and structural proteins within the mature cnidocytes that have fully developed capsule. We show that a significantly large number of upregulated genes and biochemical pathways within cnidocytes are yet to be characterized, and thus, there is a remarkable paucity of knowledge regarding the biology of cnidocytes. This study led to the identification of numerous novel protein coding genes that show cnidocyte-specific expression. Most of them surprisingly exhibited spatiotemporal variation in expression, and indicated the presence of a large population of uncharacterized cnidocytes. We also identified multiple duplication events that led to the recruitment of a bilaterian stress response complex into cnidocytes, and we highlight how it could be significant for nematogenesis. Thus, we provide several fascinating insights into the biology, development, and the evolution of cnidocytes, which are also the first venom injecting cells to originate within animals, nearly 600 MYA.

## Materials and Methods

### Sea anemone culture

*Nematostella vectensis* polyps were grown in 16 % sea salt water at 17° C. Polyps were fed with *Artemia salina* nauplii three times a week. Spawning of gametes and fertilization were performed as previously described (Genikhovich and Technau 2009b).

### Transgenesis

For generating a transgenic line expressing memOrange2 under the native regulatory region of *NvNcol-3* we microinjected a fertilized *N. vectensis* zygote with a mixture that included guide RNA (250 ng/μl) of the sequence GCAGUAGUUAGGGCAUCCCGG, Cas9 recombinant protein with nuclear localization signal (500 ng/μl; PNA Bio, USA) and a donor plasmid that includes two homology arms (spanning positions 1,380,459-1,381,924 and 1,381,925-1,383,035 in scaffold 23 of the *N. vectensis* genome) spanning the memOrange2 gene. The expression of the transgene in injected zygotes and embryos was monitored under a Nikon SMZ18 fluorescent stereomicroscope equipped with a Nikon Ds-Qi2 monochrome camera and an Elements BR software (Nikon, Japan).

### Dissociation of tentacles and cell sorting

Tentacles of *Nematostella* polyps were dissociated using a combination of papain (3.75 mg/ml; Sigma-Aldrich: P4762), collagenase (1000U/ml; Sigma-Aldrich: C9407) and pronase (1 mg/ml; Sigma-Aldrich: P5147) in DTT (0.1M) and PBS (10 mM sodium phosphate, 150 mM NaCl, pH 7.4). The tentacles were incubated with the proteolytic cocktail at 22°C overnight along with Hoechst stain (Sigma-Aldrich) to stain the nucleus. This was followed by the dissociation of tissues into single cells by flicking the tubes gently and centrifugation at 400 × *g* for 15 minutes at 4°C. The pallet was gently reconstituted in PBS.

A small amount of the dissociated sample was subjected to microscopic examination to verify successful cell separation, while the remaining sample was used for Fluorescence-activated cell sorting in a FACSAria III (BD Biosciences, USA) equipped with a 488, 405 and 561 nm lasers and a 70 μm nozzle. Two distinct populations of cells were sorted and collected: a) Hoechst positive, memOrange2 negative; and b) Hoechst positive, memOrange2-positive cell populations. The cells were directly collected into TRIzol® LS reagent (Thermo Fisher Scientific) in 3: 1 reagent to sample ratio. The instrument was maintained at 4 °C throughout the procedure.

### RNA isolation, sequencing, and assembly

Total RNA from the positive and negative populations was isolated using the TRIzol® LS reagent according to manufacturer’s protocol. Following isolation, the RNA samples were treated with Turbo DNase (Thermo Fisher Scientific), followed by re-extraction with the Tri-Reagent to remove DNase. RNA quality was assessed on a Bioanalyzer Nanochip (Agilent, USA), and only samples with RNA Integrity Number (RIN) ≥ 8.0 were considered for sequencing (all samples except TP10: 7.5 RIN). Sample library preparation for RNA sequencing was accomplished using the Illumina TruSeq RNA library protocol (mean insert size of 150). The samples were sequenced on Illumina Nextseq 500 high output v2 platform (2 × 40 bp), which generated an average of 50 million paired-end reads per replicate. The Illumina basespace pipeline was used for de-multiplexing and filtering high quality sequencing reads. Additional quality filtering steps were performed using Trimmomatic v0.36 (Bolger et al. 2014) to remove adapters, leading and trailing low quality bases (below quality 3), very short reads (shorter than 20 bases), and reads with less than an average quality score of 20 using a sliding window of 4 bases. The quality of the preprocessed and the processed data was verified using FastQC (Andrews 2010). Raw sequencing data has been deposited to the Sequence Read Archive (SRA) at NCBI (Bioproject PRJNA391807 – will be released with the publication of this manuscript).

### Differential gene expression and transcript annotation

Reads were aligned to the indexed genome of *N. vectensis* (Putnam et al. 2007) using STAR (Dobin et al. 2013), followed by the quantification of gene expression with HTSeq-count v0.6 (Anders et al. 2015). Differential expression analyses were performed using two Bioconductor packages: DESeq v2.1 (Love et al. 2014) and edgeR v3.16 (Robinson et al. 2010). Genes identified in concert by these two methods were considered as differentially expressed. The assembled transcriptomes were functionally annotated using Blast2go v4.1 (Gotz et al. 2008) against Toxprot (Jungo et al. 2012) and Swissprot (May 16, 2014) databases.

### In situ hybridization (*ISH) and immunostaining*

Colorimetric ISH was performed as previously described (Genikhovich and Technau 2009a). Double Fluorescent ISH (dFISH) was performed also according to published protocols (Nakanishi et al. 2012; Wolenski et al. 2013) with tyramide conjugated to Dylight 488 and Dylight 594 fluorescent dyes (Thermo Fisher Scientific, USA). In ISH and FISH, embryos older than 4 days were treated with 2u/pl T1 RNAse (Thermo Fisher Scientific) after probe washing in order to reduce background. Immunostaining was performed according to a previously described protocol (Moran et al. 2012), employing a commercially-available rabbit polyclonal antibody against mCherry (Abcam, USA) that cross-reacts with memOrange2 or a mouse monoclonal antibody against FLAG (Sigma-Aldrich) and a custom-made guinea pig antibody against *NvNcol-3* (Zenkert et al. 2011), generously provided by Suat Özbek (Heidelberg University). In experiments where ISH and immunostaining were combined the ISH staining was performed with FastRed (Sigma-Aldrich). Stained embryos, larvae, and adult polyps were mounted in either Vectashield antifade medium (Vetor Laboratories, USA) or 85% glycerol and visualized with an Eclipse Ni-U microscope equipped with a DS-Ri2 camera and an Elements BR software (Nikon, Japan) or with a Leica SP5 upright confocal microscope equipped with 405, 488, 561 and 594 nm lasers (Leica, Germany).

### Phylogenetic analysis

Homologues of c-Jun and c-Fos transcription factors were retrieved using blast searches (Altschul et al. 1990) against NCBI’s non-redundant nucleotide sequence database, N *vectensis* genome (Putnam et al. 2007), the EdwardsiellaBase (Stefanik et al. 2014) and various cnidarian transcriptomes (Kitchen et al. 2015). The maximum-likelihood analysis was utilized for the reconstruction of the molecular evolutionary history of the c-Jun and c-Fos protein families. Trees were generated using PhyML 3.0 (Guindon et al. 2010), and node support was evaluated with 1,000 bootstrapping replicates. The sequence alignments used for tree reconstructions are made available as Supplemental File 1.

## Supplemental files

**Supplemental Table 1. Differentially expressed genes in positive and superpositive cell populations**

**Supplemental Fig. 1. Maturation of cnidocytes**

**Supplemental Fig. 2. Heatmap of differentially expressed genes in positive and Super-positive cell populations**

**Supplemental Fig. 3. Cnidocyte-specific markers**

**Supplemental Fig. 4. Biochemical pathways in cnidocytes**

**Supplemental Fig. 5. Sequence alignment of c-Fos proteins**

**Supplemental File 1. Sequence alignments**

## Author Contributions

KS and YM designed the experiments. KS, YCS, and YM performed experiments. KS performed bioinformatic analyses. KS, YCS and YM wrote the paper. RA, AF, and NG assisted in some of the experiments.

## Acknowledgments

K.S. was supported by a Marie Sktodowska-Curie Individual Fellowship (654294). This work was supported by Israel Science Foundation grant no. 691/14, awarded to Y.M., and by the Marie Sktodowska-Curie Fellowship awarded to KS and YM. We are thankful to Dr. William Breuer (Interdepartmental Unit of the Alexander Silberman Institute of Life Sciences) for his help with FACS experiments and to Dr. Naomi Melamed-Book (Imaging Unit of the Alexander Silberman Institute of Life Sciences) for her help with confocal microscopy. We are also thankful to Dr. Robert Zimmermann (University of Vienna) for his advice and support in bioinformatic analyses.

